# Precise and efficient C-to-U RNA Base Editing with SNAP-CDAR-S

**DOI:** 10.1101/2023.02.17.528953

**Authors:** Ngadhnjim Latifi, Aline Maria Mack, Irem Tellioglu, Salvatore Di Giorgio, Thorsten Stafforst

## Abstract

Site-directed RNA base editing enables the transient and dosable change of genetic information and represents a recent strategy to manipulate cellular processes, paving ways to novel therapeutic modalities. While tools to introduce adenosine-to-inosine changes have been explored quite intensively, the engineering of precise and programmable tools for cytidine-to-uridine editing is somewhat lacking behind. Here we demonstrate that the cytidine deaminase domain evolved from the ADAR2 adenosine deaminase, taken from the RESCUE-S tool, provides very efficient and highly programmable editing when changing the RNA targeting mechanism from Cas13-based to SNAP-tag-based. Optimization of the guide RNA chemistry further allowed to dramatically improve editing yields in the difficult-to-edit 5’-C<u>C</u>N sequence context thus improving the substrate scope of the tool. Regarding editing efficiency, SNAP-CDAR-S outcompeted the RESCUE-S tool clearly on all tested targets, and was highly superior in perturbing the β-catenin pathway. NGS analysis showed similar, moderate global off-target A-to-I and C-to-U editing for both tools.

## Introduction

Cytidine (C) deamination yielding uridine (U) is a well-known posttranscriptional reaction that diversifies genetic information at the RNA level.^1^ The enzymatic base conversion is carried out by hydrolases/deaminases belonging to the class of AID/APOBEC proteins, of which some are specific for RNA, while others can use both RNA and DNA, or only DNA as substrates. The first C-to-U RNA editing enzyme described was APOBEC1 (APO1),^2^ which catalyzes the switch from the long ApoB100 to the short ApoB48 isoform by rewriting a glutamine codon (5’-<u>C</u>AA) into a STOP codon (5’-<u>U</u>AA) in the enterocytes of the small intestine.^3^ Later, single-strand RNA editing activity of further members of the APOBEC family, including APOBEC3A and 3G, was discovered. However, the biological function and targets of their RNA editing activity remained unclear to some extent.^1^ C-to-U RNA editing activity is typically found only in a subset of tissues, like small intestine and liver for APO1, or monocytes and macrophages for A3A, but can be up- and down-regulated in various pathologic situations and play a role in tumor evolution,^4^ for example.^5^ AID/APOBEC enzymes are often recruited to their targets by the help of auxiliary proteins, e.g. RBM47^6^ and A1CF^7^ for APO1, and have a very strong and thus limiting preference for specific di-nucleotides as editing substrates.^1^ Highly edited substrates, like the Glutamine-to-STOP site in the ApoB transcript, are placed in specific secondary structures that assist the recruitment and activity of the deaminase.^8,9^

Targeted RNA base editing aims at harnessing C-to-U and A(denosine)-to-I(nosine) editing activity for the rewriting of genetic information, including the substitution of amino acids and the formation (C-to-U) or removal (A-to-I) of premature STOP codons.^10^ The approach opens novel avenues for drug discovery, promising to bypass technical and ethical issues related to genome editing.^10^ In this field, our lab contributed an RNA-targeting platform based on fusion proteins of the self-labeling SNAP-tag^11^ (Fig. 1a). ^12^ Initially, we engineered a programmable A-to-I RNA base editor by fusing the SNAP-tag^11^ with the catalytic domain of the RNA editing enzyme ADAR (adenosine deaminase acting on RNA).^13,14^ In these fusions, the SNAP-tag exploits its self-labeling activity to covalently tether to a guideRNA in a defined 1:1 stoichiometry by recognizing a benzylguanine (BG) moiety, the so-called self-labeling moiety, at the guideRNA. According to simple Watson-Crick base-pairing rules, the guideRNA addresses the editing of one specific adenosine residue in a selected transcript with high efficiency, broad codon scope, and very good precision.^14^ Competing RNA-targeting platforms have been developed based on Cas proteins^15^, or trans-tethering approaches.^16^ While each approach has its specific strength and weakness,^10,12^ a clear advantage of the SNAP-tag approach is its human origin, its small size, the ease of transfecting of the chemically stabilized guideRNA(s),^14^ the possibility of concurrent editing,^14^ and the ready inclusion of small molecule^17^ and photo control^18,19^. Furthermore, we have recently shown the concurrent and fully orthogonal usage of two independent RNA editing effectors by complementing a SNAP-tagged C-to-U editing effector with a HALO-tagged A-to-I editing tool within the same cell.^20^ In the latter study, we exploited the C-to-U deaminase domain from murine APO1. Other labs have developed C-to-U RNA base editing effectors based on human APO1 or APO3A. In the first example, RNA-targeting was based on the trans-tethering approach with the MS2/MCP system.^21^ In the latter, the dCas13 platform was applied.^22^ However, none of these approaches is yet working optimally. Our approach with SNAP-tagged APO1^20^ gave low editing yields on endogenous targets and its programmability, which is the addressing of any given target cytidine with a guide RNA, was somehow limited due to the strong requirement^8^ for APO1 substrates to be located in specific secondary structures. While this was better solved for the A3A target,^22^ this tool suffers from the strong substrate codon preference for 5’-U<u>C</u>. An exciting alternative came from the engineering of an artificial C-to-U editing enzyme. Specifically, laboratory evolution was used to engineer the A-to-I deaminase domain of the hyperactive E488Q mutant^23^ of the ancestor ADAR2 into a C-to-U editing enzyme.^24^ With a dCas13-based RNA-targeting mechanism the so-called RESCUE tool was steered to its target RNAs. While the programmability was good, the editing yields remained moderate for 5’-W<u>C</u> (W = A or U) codons and low for 5’-C<u>C</u>, whereas 5’-G<u>C</u> codons were hardly editable,^24^ mirroring the well-known codon preference^25^ of ADAR2. Furthermore, the RESCUE tool retained notable A-to-I off-target editing beside C-to-U off-target editing. A high-fidelity variant, RESCUE-S, was developed,^23^ that carried an additional point mutation. However, the point mutation lowered both, the C-to-U on-target and the A-to-I off-target editing yields.

**Figure 1.**
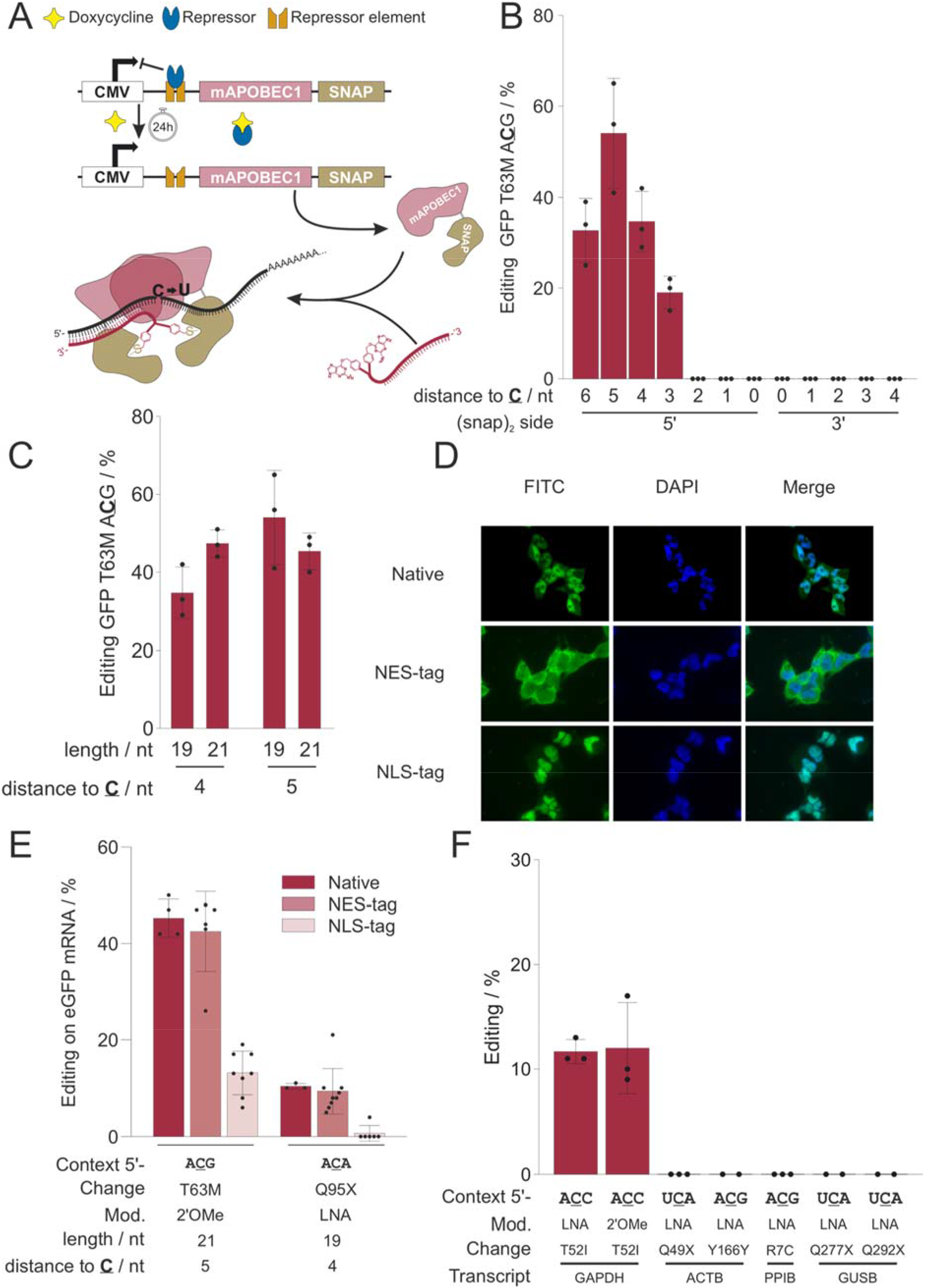
Properties of the APO1S tool. **a)** Scheme of the doxycycline-inducible APO1S tool and the guide RNA-dependent editing reaction. **b)** Dependency of editing yield of the T63M site (5’-A<u>C</u>G) in the eGFP reporter gene on the positioning of the guide RNA (19 nucleotides length, fully 2’OMe modified) relative to the target site (“**<u>C</u>**”). **c)** Fine-tuning of guide RNA length (19, 21 nt) and positioning (4 or 5 nt 5’ to the target cytidine) for optimal editing performance. **d)** Analysis of APO1S transgene expression and localization by fluorescence microscopy. The native mApobec1 sequence was either amended with an NES or NLS tag, as indicated, leading to cytosolic or nuclear localization of the APO1S protein, which was stained with SNAP-tag-reactive BG-FITC (green channel). **e)** Effect of editase localization on editing of two sites in an eGFP reporter with the respective best performing guideRNA design. **f)** Editing performance of guide RNAs (21 nt, position 5) with different backbone chemistries (2’-OMe, LNA) inducing the indicated amino acid changes at the respective endogenous transcripts demonstrating limited programmability of the tool. Data in b), c), e) and f) are shown as the mean ± s.d. of N ≥ 2 independent experiments as represented by individual data points.

Here, we now show that the high-fidelity cytidine deaminase acting on RNA (CDAR) domain from the RESCUE-S tool works very well when we replace the dCas13 domain with a SNAP-tag for RNA targeting. In particular on endogenous transcripts, the SNAP-CDAR-S outcompetes the RESCUE-S tool clearly and achieves moderate to good editing yields for all 5’-H<u>C</u> codons (H = C, A, U) under very good control of A-to-I and C-to-U bystander editing.

## Results

### The APOBEC1-SNAP tool suffers from low C-to-U editing yields and limited programmability

Recently, we demonstrated the harnessing of the murine APOBEC1 deaminase for site-directed C-to-U RNA base editing. For this, the SNAP-tag was fused to the C-terminus of the full length APOBEC1 enzyme, resulting in an editor called APO1S (Fig. 1 a).^20^ In transgenic cell lines that co-express APO1S together with SNAP-ADAR1Q, moderate editing yields were achieved on an eGFP reporter gene, but editing yields on the endogenous GAPDH transcript stayed below 20%. By targeting the eGFP reporter, we now tried several means to improve editing yields. On the guide RNA side, the positioning of the guide RNA four to five nucleotides upstream with respect to the target cytidine was most important (Fig. 1 b, c). On the protein side, the localization of the editing enzyme to the cytosol was particularly necessary (Fig. 1 d, e, Supplementary Figures S1-S6). Nevertheless, the APO1S editor suffered overall from low editing yields on endogenous targets and from low programmability (Fig. 1f), meaning that transfer to endogenous transcript was particularly difficult followed by notable guide RNA-dependent bystander editing when on-target editing was successful. (Supplementary Fig. S7).

### The SNAP-CDAR-S tool combines high editing yields with excellent programmability

In contrast, the Cas-13-mediated C-to-U editing tool called RESCUE applies a C-to-U deaminase that was evolved from the A-to-I deaminase ADAR2,^24^ and shares with ADAR2 its strong substrate preference for double-stranded RNA. Indeed, the RESCUE tool seems to have much better programmability and on-target editing was reliably obtained when the target site was positioned inside the guide RNA / mRNA duplex. However, Cas13-mediated C-to-U editing suffers from global and local C-to-U and A-to-I off-target editing, and attempts to create more precise tools, like Cas13-RESCUE-S, came along with largely reduced on target editing yields, hardly above 10% on endogenous transcripts.^24^ However, we were wondering how the engineered cytidine deaminase acting on RNA (CDAR) domain would work in the context of a SNAP-tagged fusion protein.^12^ For this, we fused the evolved, high-fidelity deaminase of the RESCUE-S tool to the C-terminus of a SNAP-tag^11,14^ to obtain the SNAP-CDAR-S tool (Fig. 2 a). We stably integrated a single copy of the SNAP-CDAR-S transgene into HEK 293 cells by using the Flp-In approach,^14,19^ and found homogenous transgene expression under control of doxycycline. Similar to the closely related A-to-I editing enzyme SNAP-ADAR2Q (Supplementary Fig. S1), the SNAP-CDAR-S tool was mainly localized to the cytosol (Fig. 2 b).

**Figure 2.**
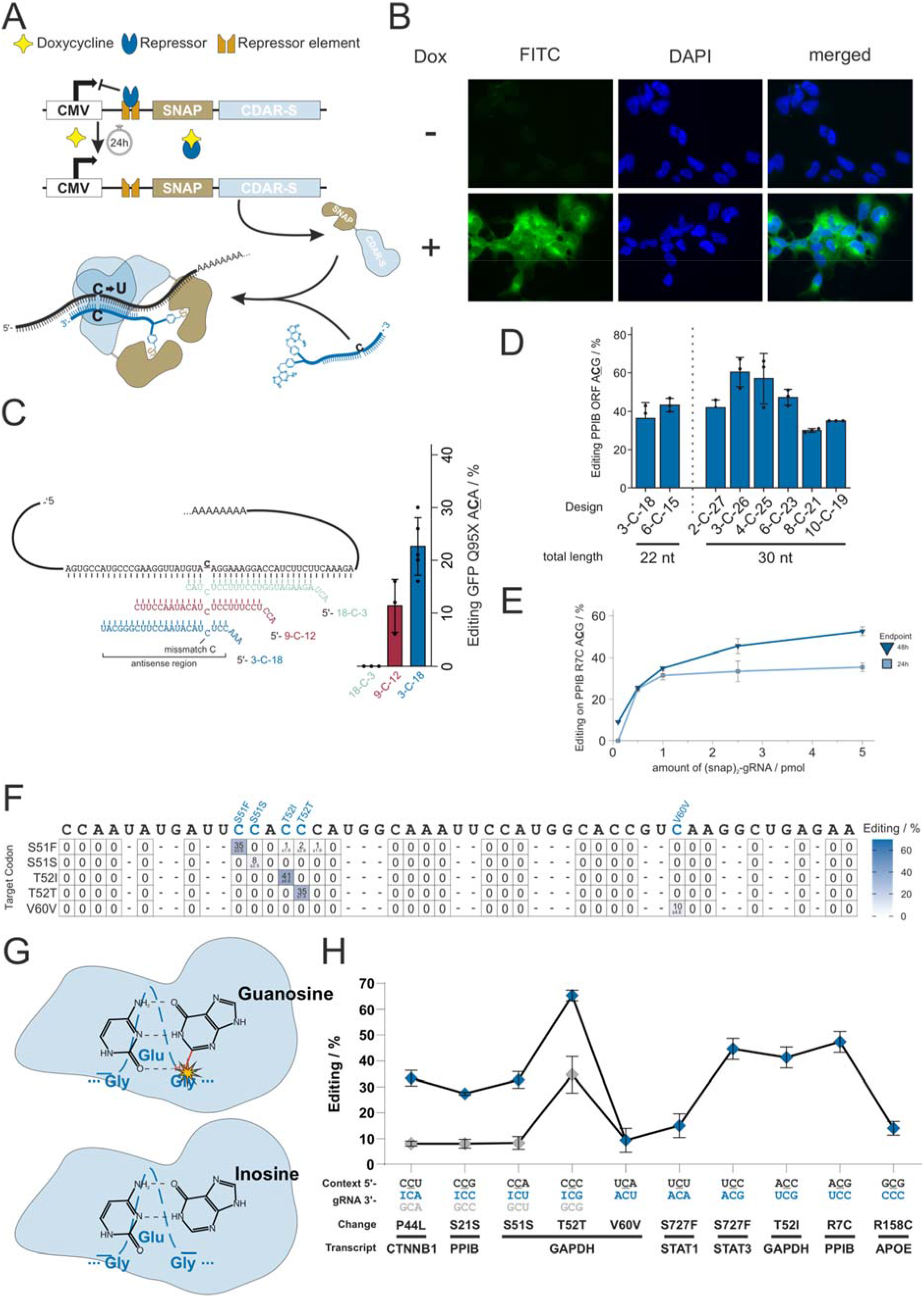
Guide RNA design and performance of the SNAP-CDAR-S tool. **a)** Scheme of the doxycycline-inducible SNAP-CDAR-S tool and the guide RNA-dependent editing reaction. **b)** Analysis of SNAP-CDAR-S transgene expression and localization by fluorescence microscopy. SNAP-CDAR-S was stained with SNAP-tag-reactive BG-FITC (green channel). **c)** Dependency of editing yield of the Q95X site (5’-A<u>C</u>A) in the eGFP reporter gene on the positioning of the guide RNA (all 22 nucleotides length, 2’OMe modification on all nucleotides except for mismatch C and flanking nucleotides) relative to the target site (symmetric versus asymmetric). **d)** Fine-tuning of guide RNA length (22, 30 nt) and positioning (of mismatch cytidine C) for optimal editing performance. **e)** Dependency of RNA editing yield on the amount of transfected guide RNA (pmol/96 well). **f)** Programmability and precision of the SNAP-CDAR-S tool. Five guide RNAs (each 6-C-23, 2’OMe gapmer design) against five nearby cytidine sites on the endogenous GAPDH transcript, each with a distinct codon context, were applied and the on-target and the C-to-U and A-to-I bystander off-target editing was determined by Sanger sequencing. **g)** Scheme explaining the positive effect of inosine in guide RNAs for targeting 5’-C<u>C</u>N codons. Pairing of the 5’ cytidine in a 5’-C<u>C</u>N context with a guanosine leads to a steric clash of CDAR’s glycine with the guanosine’s exocyclic NH_2_-group (27). Inosine lacks that NH_2_-group thus avoiding steric clash. **h)** Codon scope of the SNAP-CDAR-S tool and effect of inosine in 5’-C<u>C</u>N codons. Given are editing yields for various codons on various endogenous transcripts when applying non-optimized guide RNAs of the standard design (6-C-23, 2’OMe gapmer). For each of the four 5’-C<u>C</u>N codons (N= U, C, A, G), the editing yields of guide RNAs are compared that contained either an inosine (I) or a guanosine (G) opposite the cytidine preceding the on-target cytidine. Data in c), d), e), f), and g) are shown as the mean ± s.d. of N ≥ 2 independent experiments.

Our initial guide RNA design was inspired from our experience with the SNAP-ADAR tool and was tested for the editing of a 5’-ACA codon in a co-transfected eGFP reporter transcript. Initial guide RNAs were 22 nt long, chemically modified by 2’-O-methylation outside the target base triplet,^26^ which is the targeted cytidine plus its two closest neighboring bases, and carried a (snap)2 self-labeling moiety^20^ at the 5’-end for the recruitment of two SNAP-CDAR-S effectors per guide RNA. The exact composition, sequence and chemistry, of all guide RNAs can be found in the Supplementary Information (Supplementary Table T1a). While the SNAP-ADAR tool prefers a relatively central positioning of the target nucleobase, the SNAP-CDAR-S effector gave clearly better yields when the target cytidine was located near the 5’-terminus of the guide RNA, e.g. design 3-C-18 in Fig. 2c, in good agreement with data from the Cas13-RESCUE^24^ tool. Next, we took a closer look at the guide RNA design for the editing of the endogenous PPIB transcript, specifically, by targeting a 5’-A<u>C</u>G codon in its coding region (ORF). Here, we varied the length of the guide RNA (22 nt and 30 nt) and the positioning of the guide RNA relative to the target cytidine, see Fig. 2d. With 60% editing yield, we found the best performing guide RNA to be 30 nt long, positioning the targeted cytidine close to the 5’-end (position 4, 3-C-26) of the guide RNA in the substrate duplex. However, also other guide RNA designs gave good editing yields and the optimal design might vary to some extend for each target sequence, as suggested for the original Cas13-RESCUE-S tool.^24^ As we sought to determine a universal guideRNA design, we compared editing yields on further endogenous transcripts putting the target <u>C</u> in position 4 (3-C-26) or position 7 (6-C-23, Fig. S8). In contrast to the target site on PPIB, positioning target <u>C</u> at position 7 showed substantially higher editing yields for all selected sites (Fig. S8). In addition, we transferred designs previously reported^24^ ideal for four endogenous sites to our SNAP-CDAR-S tool and compared them to our 6-C-23 standard designs (Fig. S9). For all targets, our standard design gave similar or even better editing yield then the previously described ideal design. This indicated that 6-C-23 can be considered a universal design for the SNAP-CDAR-S tool. We then continued to analyze the performance further. Editing yields were saturating when ≥2.5 pmol / 96 well (20 nM) guide RNA were transfected (Fig. 2e). To characterize the scope, programmability and precision of the SNAP-CDAR-S tool, we targeted five different guide RNAs (all 30 nt, all 6-C-23, 2’-OMe) against five different cytidine bases, which were all located in close proximity in the ORF of the endogenous GAPDH transcript and determined on-target as well as C-to-U and A-to-I bystander editing yields. We found excellent programmability, with good on-target yields (8% to 41%) and with lacking bystander editing (detection limit Sanger sequencing ca. 5%) at neighboring cytidine or adenosine bases (Fig. 2f, the same was found for an alternative 3-C-18 guide RNA design, see Supplementary Fig. S10). 2’-O-Methylation was shown in the past to block bystander A-to-I editing very efficiently in SNAP-ADAR tools,^14,26^ and this may contribute here to the high precision of the targeted editing too. However, we were not fully satisfied with the editing yield at the 5’-C<u>C</u>A codon (GAPDH S51S), which achieved only 8% with the best design (6-C-23, a 3-C-18 design gave even <5%). This limited scope was also reported for the Cas13-based RESCUE tool^24^ and resembles the codon preference of the ancestor ADAR2^25^ protein. A recent structural analysis of the ADAR2 deaminase bound to a dsRNA substrate revealed a steric clash between the peptide backbone of glycine 489 and the minor groove face of G=C base pairs residing at the 5’ neighboring position to the target adenosine.^27^ This steric clash could be relaxed by replacing the 5’-neighboring G=C base pair with sterically less demanding I=C base pair (lacking an exocyclic amino group), simply by pairing the 5’-C<u>C</u>A target codon with a 5’-UCI sequence in the guide RNA (Fig. 2g). Indeed, we found a 3-fold improved editing yield of 32% for the respective site in GAPDH (Fig. 2H, Supplementary Fig. S10B). We then systematically tested the principle for all four potential 5’-C<u>C</u>N codons (N = A, U, G, C) and found that an inosine base opposite the 5’-neighboring cytidine always improved editing at the targeted cytidine base (Fig. 2h). Even for the well-edited 5’-C<u>C</u>C codon (34%), we could still achieve a notable gain in editing yield (66%).

### SNAP-CDAR-S clearly outperforms Cas13-RESCUE-S on endogenous targets

To benchmark the SNAP-tagged tool with the Cas13-based tool, we generated an analogous 293 Flp-In T-REx cell line stably expressing the Cas13-based RESCUE-S on doxycycline induction. In the original work,^24^ RESCUE-S has always been applied by means of transient overexpression, however, this often leads to high variability in editing yields and artifacts in off-target analyses.^12^ We tested both editing tools side-by-side for the editing of eight different sites on five different endogenous transcripts (GAPDH, PPIB, CTNNB1, STAT3, STAT1) and one disease-relevant cDNA (APOE). Most target sites were taken from the original Cas13-RESCUE-S publication^24^ so that optimal Cas guide RNAs have already been reported for each of them (Supplementary Table T 1b). We repeated these experiments by transfecting 300 ng/96 well of the plasmid-borne optimal guide RNAs into the stable Cas13-RESCUE-S cells. The guide RNAs for the SNAP-CDAR-S cell lines were not optimized, but we simply transfected 5 pmol/96 well chemically stabilized, 30 nt guide RNAs of the 6-C-23 standard design. Nevertheless, the SNAP-CDAR-S tool clearly outcompeted the Cas13-RESCUE-S tool on all eight targets, achieving editing yields between 10% and 50% (Fig. 3a), while the editing yields of the Cas13-RESCUE-S tool did not achieve editing yields above 10%, in accordance with the original report^24^. For four targets, only SNAP-CDAR-S, but not Cas13-RESCUE-S, was able to achieve detectable editing (PPIB S21G, CTNNB1 H63Y and P44L, STAT1 S727F). Interestingly, three out of these four examples target 5’-C<u>C</u>N codons, which is readily done by the SNAP-CDAR-S approach with inosine-containing guide RNAs, highlighting that the SNAP-CDAR-S approach does not only give generally higher editing yields but also increases the codon scope towards 5’-C<u>C</u>N sites.

**Figure 3.**
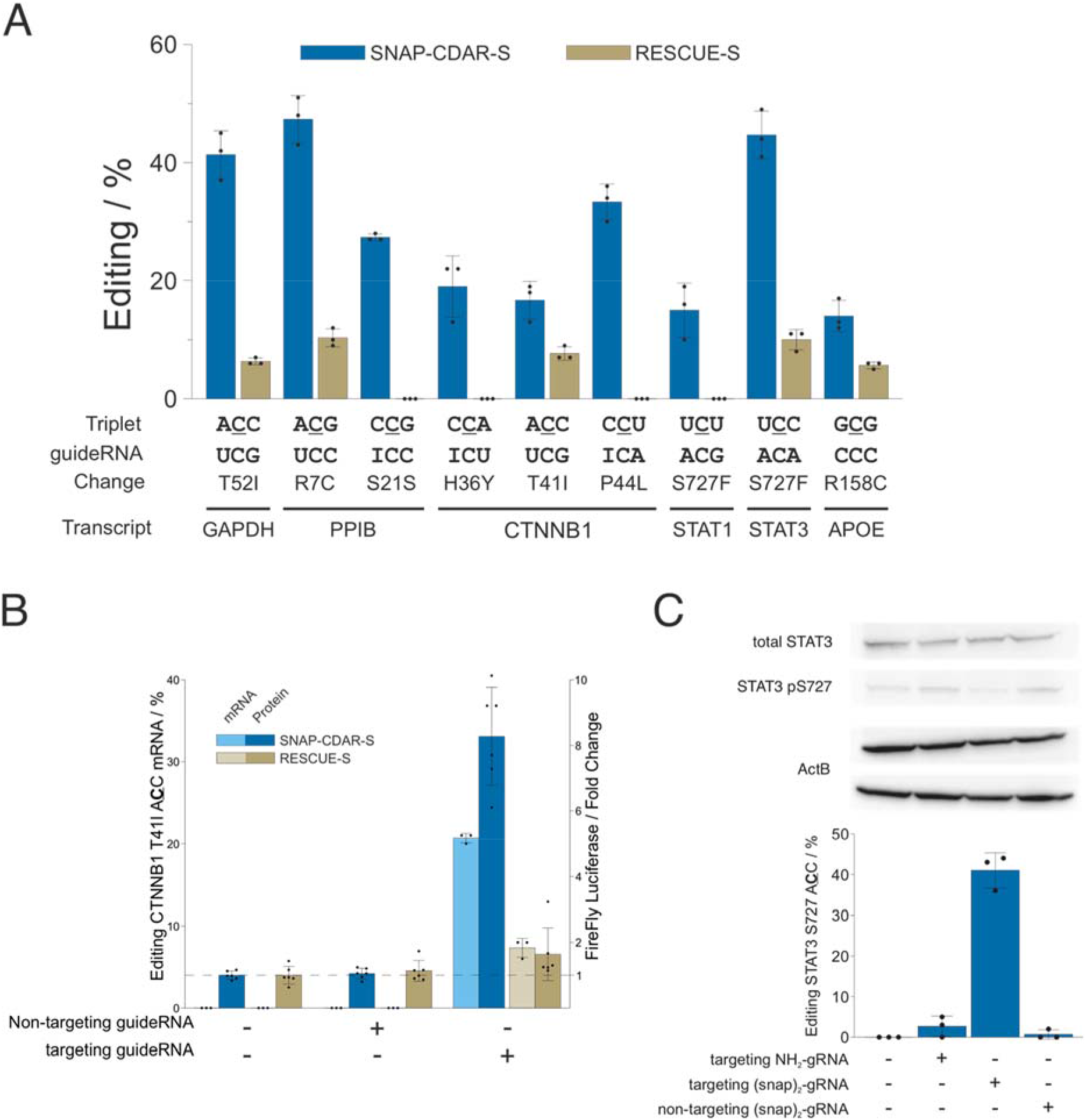
Benchmark with Cas13-RESCUE-S on endogenous targets and applications. **a)** Comparison of editing yields at various sites on various endogenous targets and one disease-relevant cDNA (APOE) comparing SNAP-CDAR-S with standard guide RNAs (30 nt, 6-C-23, 2’OMe gapmer) versus Cas13 RESCUE-S with plasmid-borne optimized Cas guide RNAs. Both editing enzymes were expressed from the same single genomic locus. **b)** Comparing both tools, SNAP-CADR-S versus Cas RESCUE-S, for the activation of β-catenin by RNA editing. Given are C-to-U editing yields (T41I) and the luminescencebased read-out of pathway activation. For further controls, see Supplementary Fig. S11. **c)** Editing of the regulatory phospho-site serine 727-to-glycine in STAT3, read-out of editing yield by Sanger sequencing, and amount of total STAT3 and pS727 STAT3 protein by Western blot. Data in a), b), and c) are shown as the mean ± s.d. of N = 3 independent experiments.

Several of the indicated targets are of clinical interest. The removal of threonine 41 from the β-catenin protein inactivates a degron and thus stabilizes the protein.^28^ Enhanced levels of β-catenin could be applied to boost liver regeneration or wound healing transiently.^29^ We benchmarked the activation of the Wnt pathway by the SNAP-CDAR-S versus its analog Cas13 tool in a plasmid-borne luciferase assay, following a protocol reported before^24^. While the SNAP-tagged tool achieved 21% editing yield and an 8-fold increase in β-catenin activity (Fig. 3b, Supplementary Figure S11), the Cas13-RESCUE-S tool gave only 7% editing yield and 1.6-fold increase. The STAT3 protein (Signal transducer and activator of transcription 3) is a multifunctional signaling molecule, which acts as a transcription factor in the nucleus, or translocates to the mitochondrium, and modulates immune response and metabolism. Its hyperactivation plays an important role in autoimmune disease, sterile inflammation, and cancer.^30^ Here, we removed serine 727 from STAT3, a functionally important phosporylation site. We could achieve up to 41% serine-to-glycine editing, which was accompanied by a visible reduction in S727 phosphorylation as determined by Western blot (Fig. 3c, Supplementary Figure S12). Finally, we aimed at introducing a protective genotype into the apolipoprotein E (APOE) transcript, introducing the rs7412 SNP (R158C), which could transfer the neutral ε3 allel (ca. 78% caucasian carriers) into the protective ε2 allel, which was shown to largely reduce the risk for atherosclerosis.^31^ However, given the non-preferred nature of the codon (5’-G<u>C</u>G), and the very high GC content of the surrounding sequence space, an editing yield of only 14% was achieved, still clearly outcompeting the Cas13 tool (Fig. 3a).

### Both tools show moderate global A-to-I and C-to-U off-target editing

We used next-generation sequencing of the poly(A)+ transcriptome (10 GB per condition) to assess the transcriptome-wide A-to-I and C-to-U off-target editing of the SNAP-CDAR-S versus its analog Cas13-RESCUE-S. We took RNA from cells expressing the respective editing effector in the presence and absence of the respective guide RNA and compared them to Flp-In T-REx cells not expressing an engineered effector^14^. First, we compared the editing reactions against the empty Flp-In T-REx cell line and were able to detect the on-target editing event (PPIB Arg7Cys) with editing yields of 48% for SNAP-CDAR-S and 14% for Cas13-RESCUE-S, which were confirmed by Sanger sequencing (Fig. 4a). Beside the on-target editing, we found around 1000 A-to-I and three to four hundred C-to-U off-target events for both effectors (Fig. 4b). As seen for ADAR-based effectors before,^10,12,32^ A-to-I off-target editing was a combination of enhanced editing at known sites and editing at novel sites, whereas the large majority of C-to-U off-target editing were novel sites. We further analyzed the outcomes of the editing reactions and found that only a moderate number of all editing events resulted in missense mutations (Fig. 4c). At less than ten missense sites, the change in the off-target editing yield was increased above 25%, indicating that most missense sites are only marginally affected (Fig. 4d). The patterns between the two effectors were very similar, which was expected given that the CDAR domain of both editing tools is identical. The presence of the editing tools gave no larger changes in gene expression (Supplementary Fig. S13), and both editing effectors were expressed to similar TPM levels (Fig. 4e). Finally, we analyzed the guide RNA-dependent changes in editing. Clearly, the vast majority of off-target editing came from the presence of the editing enzymes and was guide RNA-independent (Fig. 4f,g). The guide RNA-dependent C-to-U editing was very clean for the SNAP-CDAR-S tool. In contrast, the on-target editing with Cas13-RESCUE-S was covered by a small number of further editing events. This became also visible when we plotted all guide RNA-dependent changes in editing yields (Fig. 4h). While the on-target site gave the largest ΔEditing value for the SNAP-CDAR-S tool, the Cas13-RESCUE-S tool gave six A-to-I and another six C-to-U off-target events with higher change in editing level. Overall, both enzymes were expressed to comparable TPM levels, gave very similar patterns and levels of global off-target editing and mainly differed in the 5-fold higher on-target editing yield of the SNAP-CDAR-S tool.

**Figure 4.**
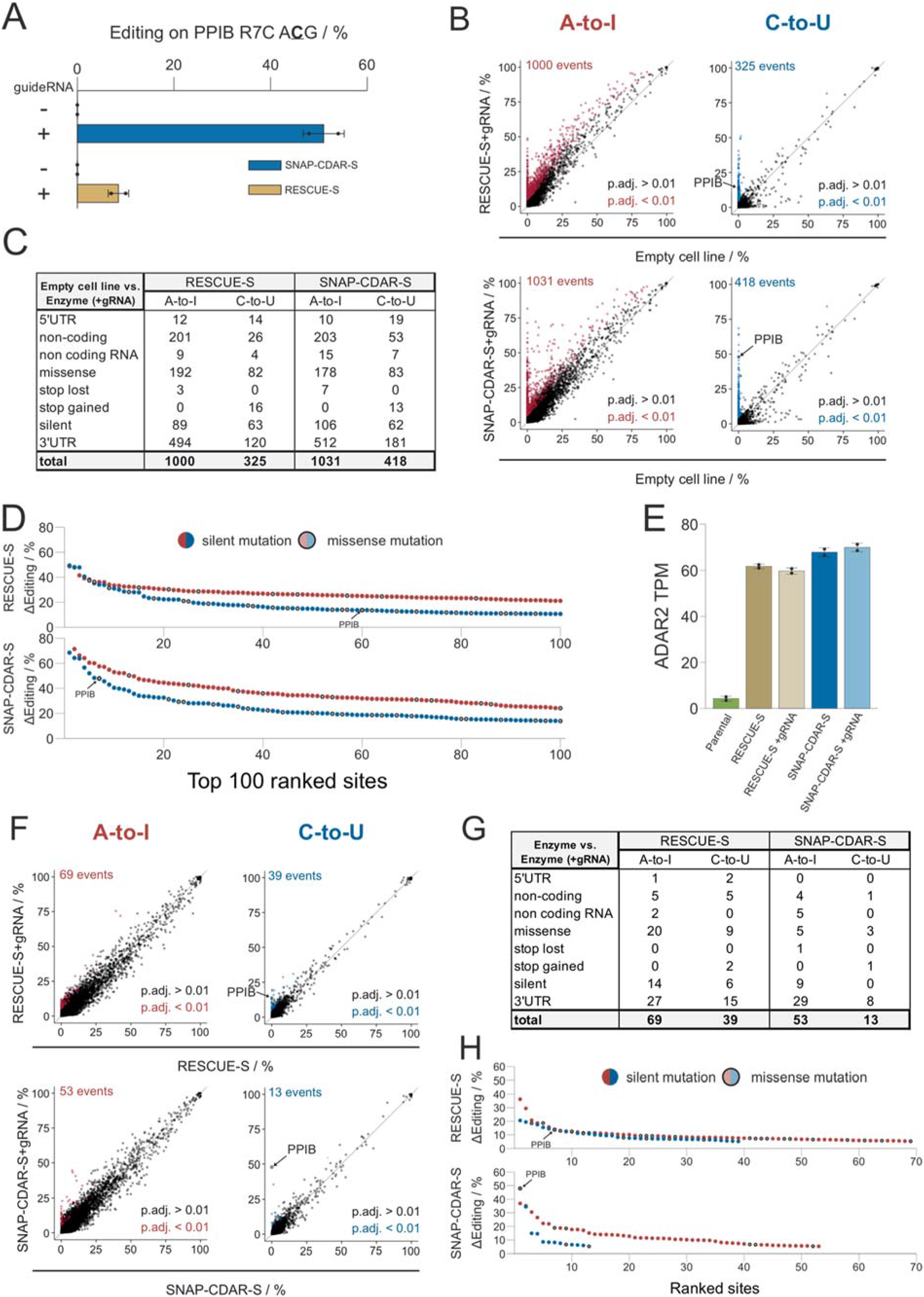
Precision of SNAP-CDAR-S versus Cas13-RESCUE-S as determined by NGS. **a)** Editing yields as determined by Sanger sequencing prior to NGS. **b)** Plot of total off-target events of the respective effector + guide RNA against the empty cell line. Significantly differently edited sites (adjusted p-value <0.01) are colored in red (A-to-I) and blue (C-to-U), respectively. **c)** Total off-target effects sorted by categories. **d)** Changes in editing yields (ΔEditing) of the 100 top-ranked editing sites color-coded for A-to-I (red) and C-to-U (blue) editing. The on-target editing event is marked by an arrow. Missense and silent mutations are indicated by empty and filled symbols, respectively. **e)** Transgene expression levels as determined by TPM values of ADAR2 (ADARB1). Given that CDAR-S is an ADAR2 deaminase mutant, both transgenes (SNAP-CDAR-S, Cas13-RESCUE-S) were annotated as such. ADAR2 itself is not expressed in the parental 293 cell line. Both transgenes were expressed to comparable TPM levels and did not change upon presence of the guide RNA. **f)** Plot of guide RNA-dependent off-target events of the respective effector + guide RNA against the respective effector. Significantly differently edited sites (adjusted p-value <0.01) are colored in red (A-to-I) and blue (C-to-U), respectively. **g)** Guide RNA-dependent off-target effects sorted by categories. **h)** Changes in editing yields (ΔEditing) of the topranked editing sites, color-coded for A-to-I (red) and C-to-U (blue) editing. The on-target editing event is marked by an arrow. Missense and silent mutations are indicated by empty and filled symbols, respectively. Significance in panel b) and f) was tested by Fisher’s exact test (two-sided), n = 2 independent experiments.

## Discussion

In comparison to the APOBEC1 enzyme, the CDAR domain, evolved from ADAR2, performs considerably better in targeted RNA base editing tools. While the CDAR domain was taken from the Cas13-mediated RESCUE approach,^24^ we could show here that this deaminase domain works particularly well when the self-labeling SNAP-tag is applied as the RNA-targeting mechanism. Compared to the Cas13-RESCUE-S, the SNAP-CDAR-S gave reliably higher on-target yields with less bystander editing, while the global A-to-I and C-to-U off-target effects were comparable. A reason for the superior efficiency of the SNAP-tagged tool might be the chemical design and the covalent bond that tethers the guide RNA to the SNAP-tag and may foster the encounter of guide RNA, target RNA and editing effector.^12,14^ Regarding the design of the guide RNAs with respect to chemical modifications, we found that lessons learned from the engineering of the closely related SNAP-ADAR tool^14,26^ could be largely transferred. In particular, the general guide RNA design with 2’-O-methylation at the ribose moieties outside the base triplet and the usage of non-encodable bases like inosine opposite 5’-C<u>C</u>N codons have contributed to the improved performance so that editing yields between 10% and 50% are regularly achieved in 5’-H<u>C</u>N (H = C, A, U) codons, and only 5’-G<u>C</u>N codons remain challenging. To our knowledge, our data is the first report of stable integration of the Cas13-RESCUE-S tool and shows that it functions as well under genomic integration as it does via plasmid overexpression. Even though Cas13-RESCUE-S was presented as a high-fidelity enzyme with largely reduced global A-to-I off-target editing before,^24^ there still remains notable A-to-I as well as C-to-U off-target editing, which might have been underestimated in the prior study, where off-target analysis was done on reporter cDNA under co-transfection of the editing enzyme. Our work further allows to compare the A-to-I off-target effects of the SNAP-CDAR-S deaminase directly to the related, stably integrated wildtype and hyperactive (E488Q) mutant of SNAP-ADAR2^14^. While the off-target A-to-I editing of the SNAP-CDAR-S tool is clearly below that of the hyperactive, off-target-prone SNAP-ADAR2 E488Q mutant,^14^ the SNAP-CDAR-S tool still has notably frequent off-target A-to-I editing when compared to the wildtype SNAP-ADAR2 enzyme (see Supplementary Fig. S14), which is clearly not yet optimal for a C-to-U editing enzyme and may require further engineering efforts to generate a pure C-to-U editing enzyme. Overall, the SNAP-CDAR-S tool adds a reliable and efficient enzyme to the targeted RNA base editing toolbox. Even though editing efficiency and precision are not yet perfect, the tool clearly outperforms the Cas13-based RESCUE-S and lays a basis for further engineering in the future.

## Data availability

The raw NGS data will be uploaded to the GEO server (https://www.ncbi.nlm.nih.gov/geo/) prior to publication.

## Acknowledgments

We would like to express our gratitude to Dr. Nina Papavasiliou and Dr. Rafail Tasakis for their expertise and support on the analysis of the Next Generation Sequencing data.

